# Symptoms of Depression in a Large Healthy Population Cohort are related to Subjective Memory Complaints and Memory Performance in Negative Contexts

**DOI:** 10.1101/101642

**Authors:** Susanne Schweizer, Rogier A. Kievit, Tina Emery, Cam-CAN, Richard N. Henson

## Abstract

Decades of research have investigated the impact of clinical depression on memory, which has revealed biases and in some cases impairments. However, little is understood about the effects of sub-clinical symptoms of depression on memory performance in the general population. Here we report the effects of symptoms of depression on memory problems in a large population-derived cohort (N = 2544), 87% of whom reported at least one symptom of depression. Specifically, we investigate the impact of depressive symptoms on subjective memory complaints, objective memory performance on a standard neuropsychological task and, in a subsample (n = 288), objective memory in affective contexts. There was a dissociation between subjective and objective memory performance, with depressive symptoms showing a robust relationship with self-reports of memory complaints, even after adjusting for age, gender, general cognitive ability and symptoms of anxiety, but not with performance on the standardised measure of verbal memory. Contrary to our expectations, hippocampal volume (assessed in a subsample, n = 592) did not account for significant variance in subjective memory, objective memory or depressive symptoms. Nonetheless, depressive symptoms were related to poorer memory for pictures presented in negative contexts, even after adjusting for memory for pictures in neutral contexts. Thus the symptoms of depression, associated with subjective memory complaints, appear better assessed by memory performance in affective contexts, rather than standardised memory measures. We discuss the implications of these findings for understanding the impact of depressive symptoms on memory functioning in the general population.

## Symptoms of Depression in a Large Healthy Population Cohort are related to Subjective Memory Complaints and Memory Performance in Negative Contexts

Mood fluctuates throughout the day as well as the lifespan, though overall most individuals feel “fine” most of the time (Taquet et al., 2016). At one time or another, however, virtually everyone will experience low mood that is significant enough to endorse one or more symptoms of depression. In 24% of the cases, this will be severe enough to meet diagnostic criteria for major depressive disorder (Horwath et al., 1992; Kessler et al., 2005). Much needed research has been dedicated to the affective, cognitive, and neurobiological correlates of these major depressive episodes (Davidson et al., 2002; Gotlib and Joormann, 2010; Menon, 2011). Less however is known about the cognitive effects of depressive symptoms within the sub-clinical range commonly experienced by the general population. Here we explore the impact of depressive symptoms on memory, as experienced by a population cohort of over 2500 adults that was specifically selected for being currently free of neuropsychiatric disorders (the Cambridge Centre for Ageing and Neuroscience (Cam-CAN) cohort; www.cam-can.org). Importantly, these individuals were tested on a range of memory measures including subjective memory complaints, performance on a standardized measure of memory, and performance (in a subset of the cohort) on a task specifically designed to assess memory in affective contexts.

The findings from the clinical literature suggest the ability to recall relevant information over time is reduced in individuals who experience clinical levels of depression (Burt et al., 1995; Rock et al., 2014), although there is some suggestion that this may be due to more general reductions in executive functioning (Elderkin-Thompson et al., 2007; e.g., Fossati et al., 2002). One possible neurobiological mechanism for these memory problems is prolonged exposure to elevated levels of corticosteroids, owing to the heightened psychological stress experienced in a major depressive episode (Gotlib et al., 2008; Lamers et al., 2013; MacQueen and Frodl, 2011; Pariante and Lightman, 2008; Reppermund et al., 2007). The animal literature shows robust, well-replicated associations between stress exposure, levels of corticosteroids and memory performance, specifically through the neurodegenerative effect of corticosteroids on the hippocampus (Kim and Diamond, 2002; McEwen, 1999). The hippocampus is critical to the consolidation of information into long-term memory (Bird and Burgess, 2008; Neves et al., 2008). Studies in human depression yield more equivocal results, though evidence generally supports the theory of volumetric shrinkage of the hippocampal complex in individuals suffering from depression (Campbell et al., 2004; Fried and Kievit, 2016; Hastings et al., 2004; MacQueen and Frodl, 2011; Videbech and Ravnkilde, 2004). However, the field faces methodological challenges, some of which are inherent to the population under investigation. For example, comparing brain abnormalities across studies entails comparing across subtypes of depression, different levels of chronicity, environmental factors and variations in exposure to psychotropic medication (Fried et al., 2014). Moreover, little is known about how the effects of clinical depression on memory relate to the effects of (subclinical) symptoms of depression experienced in the general population. Building on the clinical literature, the present study aims to address this gap. We predicted that symptoms of depression experienced in the general population would be related to memory impairments (as assessed by a standardized measure of memory), and that this relationship would be associated with smaller hippocampal volumes.

The association between depressive symptoms and subjective memory complaints has been the focus of considerable previous research, especially in individuals in the later stages of their life (Bolla et al., 1991; Comijs et al., 2002; Flicker et al., 1993; Grut et al., 1993; Jonker et al., 2000; Jorm et al., 2001; O’Connor et al., 1990; Schmand et al., 1997). Research into the predictive utility of subjective memory complaints for objective memory performance and dementia diagnoses in older adults has revealed that subjective memory complaints are better accounted for by individuals’ levels of depressive symptomatology than their actual memory performance (Derouesné et al., 1999; O’Connor et al., 1990; Reid and MacLullich, 2006; Schofield et al., 1997; Smith et al., 1996). In line with these findings, we predicted that subjective memory complaints would increase as a function of symptoms of depression. Negative interpretative biases observed in those with clinical levels of depression, which increase as a function of symptoms of depression in non-clinical populations, may partially account for this finding (Beck, 2008; Mathews and MacLeod, 2005). Alternatively the association between self-reported symptoms of depression and self-reported memory problems may simply reflect a response tendency on measures assessing neuropsychiatric health complaints. The current sample allowed a direct test of the latter hypothesis, by using the Hospital Anxiety and Depression Scale (Olssøn et al., 2005; a questionnaire with adequate psychometric properties; HADS; Zigmond and Snaith, 1983) to assess symptoms of depression. The two subscales of the HADS assess symptoms of anxiety and depression, respectively. If increased memory complaints reflect a simple response tendency, then symptoms of depression and anxiety should show the same association with subjective memory complaints. In contrast, if the increase in self-reported memory complaints is specific to depressive symptomatology, then subjective memory complaints should be more reliably associated with symptoms of depression than symptoms of anxiety.

Previous work additionally suggests that altered cognitive and affective processing in depression are associated with other changes in memory performance. For example, depressed individuals exhibit a mood-congruency bias, which makes them able to recall more negative memoranda compared to non-depressed individuals (Elliott et al., 2002). Another memory phenomenon observed in individuals with depression is that their autobiographical memories lack specificity (e.g., when asked to recall an instance of a specific birthday party, a depressed individual might respond ‘my father was never present at my birthday parties’, 42). These deviations from typical memory performance suggest an abnormality in basic memory operation and/or in the processing of affective information. Research on memory for affective stimuli and events more broadly shows that affective information is better remembered than neutral information (LaBar and Cabeza, 2006). What remains under-researched is the effect of affective context on memory performance. Henson and colleagues (Henson et al., 2016) have recently shown that, in the same CamCAN cohort as is studied here, recognition memory for neutral objects varied as a function of the affective valence (negative, positive, or neutral) of the background context against which those objects were originally presented. The increased affective significance (cf. 2009) of negative information to individuals who currently experience symptoms of depression is likely to attract attentional resources towards negative backgrounds and away from neutral objects superimposed on those backgrounds, thereby impairing the encoding into memory of those objects. For this reason, memory for information presented in affective contexts may be more sensitive to the influence of subclinical depressive symptoms than the more commonly-used, affect-neutral measures of memory.

In summary, the present study investigated the hypotheses that depressive symptoms are related to more subjective memory complaints (*Hypothesis 1a*) and worse objective memory performance (*Hypothesis 1b*). This first pair of hypotheses was investigated in all individuals from the CamCAN cohort who completed all measures of interest during an interview assessment in participants’ homes (*N* = 2544). The study further explored whether the relationship between memory performance and depressive symptoms is related to reductions in hippocampal volumes (*Hypothesis 2*). This was investigated in a subsample (*n* = 592) for whom volumetric data of the hippocampus were available from a more extensive neurocognitive assessment including the acquisition of T1- and T2-weighted MRI scans. The third prediction was that self-reported symptoms of depression would be more strongly related to a measure of memory in negative contexts compared to a standard measure of memory (*Hypothesis 3*). This hypothesis was investigated in a second subsample (*n* = 288) that completed a more specialized memory task.

The nature of the study’s sample also allowed for a number of additional explorations: first, as outlined above, we tested whether H1a was specific to symptoms of depression. That is, whether the relationship between depressive symptoms and self-reported memory complaints reflected a general response tendency toward reporting more neuropsychiatric complaints and would therefore show the same relationship with symptoms of anxiety. Next, given the evidence suggesting that memory problems related to depressive symptoms may be a function of general impairments in cognitive ability observed in individuals experiencing symptoms of depression (Elderkin-Thompson et al., 2007; Fossati et al., 2002), the study investigated whether symptoms of depression remained significantly related to the various types of memory after controlling for general cognitive ability. Third, the study explored whether the relationships in hypotheses 1-3 would remain after accounting for variations in memory, hippocampal volume and depressive symptoms attributable to age (Ferrari et al., 2013; Grady and Craik, 2000; Hedden and Gabrieli, 2004; Jorm, 2000). And finally, because women tend to show better verbal recall performance and more symptoms of depression, we investigated whether gender differences contribute to the relationships predicted in hypotheses 1-3 (Andreano and Cahill, 2009; Piccinelli and Wilkinson, 2000).

## Methods

### Participants

The full sample included 2544 individuals from the CC3000 Cam-CAN sample (Shafto et al., 2014). These participants (95% of the total Cam-CAN sample) were included because they had completed all measures pertaining to our first hypothesis. Structural imaging data was available for 592 participants from our overall sample. Hypothesis 2 was tested on this subsample. Finally, 288 participants from the total sample completed the valenced memory task and were included in the investigation of our third hypothesis. See Table S1 for participant characteristics.

### Measures

#### Depressive symptoms

Symptoms of depression were assessed with the depression subscale of the HADS (Zigmond and Snaith, 1983). The subscale consists of 7 items for which participants indicate how frequently they have felt them over the past week on a scale form “0” = “Not at all” to “3” = “Most of the time”. The scales have been well validated for use in the general population (for a review see 41).

#### Objective memory

Objective memory was assessed with a standard measure of memory, the delayed recall of a story taken from the Wechsler Memory Scale (Wechsler, 1999).

#### Subjective memory

Participants were simply asked whether they experienced any memory problems or not: ‘Do you feel you have problems with your memory? Yes/No.’.

#### Valenced memory

Memory for objects in positively and negatively valenced as well as valence neutral contexts was assessed with a newly designed measure (Henson et al., 2016). The task consisted of a study and a test phase. The study phase was divided into two ten minute blocks with a short break in between. In each block participants were presented with 60 background images selected from the International Affective Picture System (Lang et al., 2008) that appeared on the screen for 2 seconds before an object was superimposed on the background image. The object and background stayed on the screen for 7.5 seconds. Participants were asked to press the button as soon as they had come up with a story to help them link the object and background together. They were asked to keep elaborating on that story until the object and background disappeared. There was a 0.5 interval second before the next trial started.

Participants were advised that some images would be pleasant and others unpleasant, but they were not informed that their memory for the items and their background would be tested later. After participants completed the second block they were given a 10 minute break before the test phase. In the test phase they saw 160 trials that were split into four 20 minute blocks. The trials included 120 studied objects and 40 new objects. Each test trial first presented participants with a masked (pixel noise) picture of an object and participants had name the object or respond “I don’t know” before pressing the key to reveal the object. Memory for the object was then tested by asking participants whether or not the object had been presented in the study phase. Participants then indicated how confident they were of their response by pressing one of four keys: “sure new”, “think new”, “sure studied”, or “think studied”. For trials on which participants indicated “studied” sure or think, their associative memory was tested. That is, they were asked to say out-loud whether the object had been presented over a positive, neutral or negative background or to respond “I don’t know”. Finally, they were asked to describe the background image. The priming, associative memory and qualitative data are not reported as part of this study.

Memory accuracy was computed as the *d’* measure of discriminability (Green and Swets, 1966): *d’* = Φ^−1^(*pH*)−Φ^−1^(*pFA*). *pH* denotes the proportion of hits, *pFA* the proportion of false alarms and Φ^−1^ the inverse cumulative distribution function of the Normal distribution (*d’* = 0 for chance performance; extreme values of 0 or 1 for *d’* were adjusted using a log-linear approach).

#### General cognitive functioning

In the overall cohort, cognitive ability was assessed with the Addenbrooke’s Cognitive Examination – Revised assessment (ACE-R) (Mioshi et al., 2006). The screening measure was devised to detect signs of dementia and cognitive impairment assessing five domains of cognitive functioning: orientation/attention, memory, verbal fluency, language and visuospatial ability. The memory domain assess both immediate and delayed recall. As with our assessment of objective memory, participants had to recall verbal information after a delay interval. We therefore used the ACE-R sum score of all domains except memory. For the neuroimaging and affective subsamples, the ACE-R was not an informative test of cognitive ability due to participants scoring at ceiling. We therefore included the Cattell’s culture-free test of intelligence (Cattell, 1971), which was available in both subsamples (not the overall cohort). The test requires participants to complete complex pattern matrices, and has previously shown strong associations with behavioral and neural domains within the Cam-CAN cohort (Kievit et al., 2014).

### Structural MRI

Gray-matter was estimated from the combined segmentation of 1mm^3^, T1- and T2-weighted MR images, followed by diffeomorphic registration of grey-matter segments from all participants in Stage 2 of the CamCAN study in order to create a sample-specific template. This template was then transformed into Montreal Neurological Institute (MNI) space, and every participant’s grey-matter image resliced to the same space, while being modulated by the warping entailed. These stages were done in SPM12 (www.fil.ion.ucl.ac.uk/spm). For details of the MRI sequences, see (Shafto et al., 2014) for further details of the MRI preprocessing, see (Taylor et al., In Press). Grey-matter volume (GMV) of Hippocampus was estimated by summing all voxel values within the left and right Hippocampal regions of interest (ROIs) from the Harvard-Oxford Atlas (http://fsl.fmrib.ox.ac.uk/fsl/fslwiki/Atlases).

### Procedure

After providing informed written consent, participants completed the “Stage 1 – Interview” of the Cam-CAN study including computerized health and lifestyle questionnaires as well as a core cognitive assessment (Shafto et al., 2014). The subset of measures included in the present study are described below. A subset of the sample (subsamples selected to investigate hypotheses 2 and 3, see above) were included in the “Stage 2 – Core Cognitive Neuroscience” phase of the Cam-CAN study. As part of the second stage of the study, participants completed a series of cognitive task across three sessions. One of these sessions also included core structural and functional MRI measures. This second stage also included the valenced memory task. Given the time constraints across multiple sessions, the Cam-CAN protocol included a subset of tasks, including the valenced memory tasks, administered to only a randomized subset (50%) of participants. This study complied with the Helsinki Declaration, and was approved by the Cambridgeshire 2 Research Ethics Committee (reference: 10/H0308/50).

## Results

Figure 1 shows the distribution of scores from the depression sub-scale of the HADS, while Table S1 provides a comprehensive overview of the characteristics of all three samples. We found that 87% (*n* = 2211) of participants reported at least 1 symptom of depression, and over 91% scored a total of 7 points or less on the depression sub-scale (Zigmond and Snaith, 1983), which is below the commonly reported clinical cutoff of 8 points for this sub-scale (Bjelland et al., 2002). This shows that symptoms of depression are common in the general population, but typically do not reach clinically significant levels^1^.

**Figure 1.**
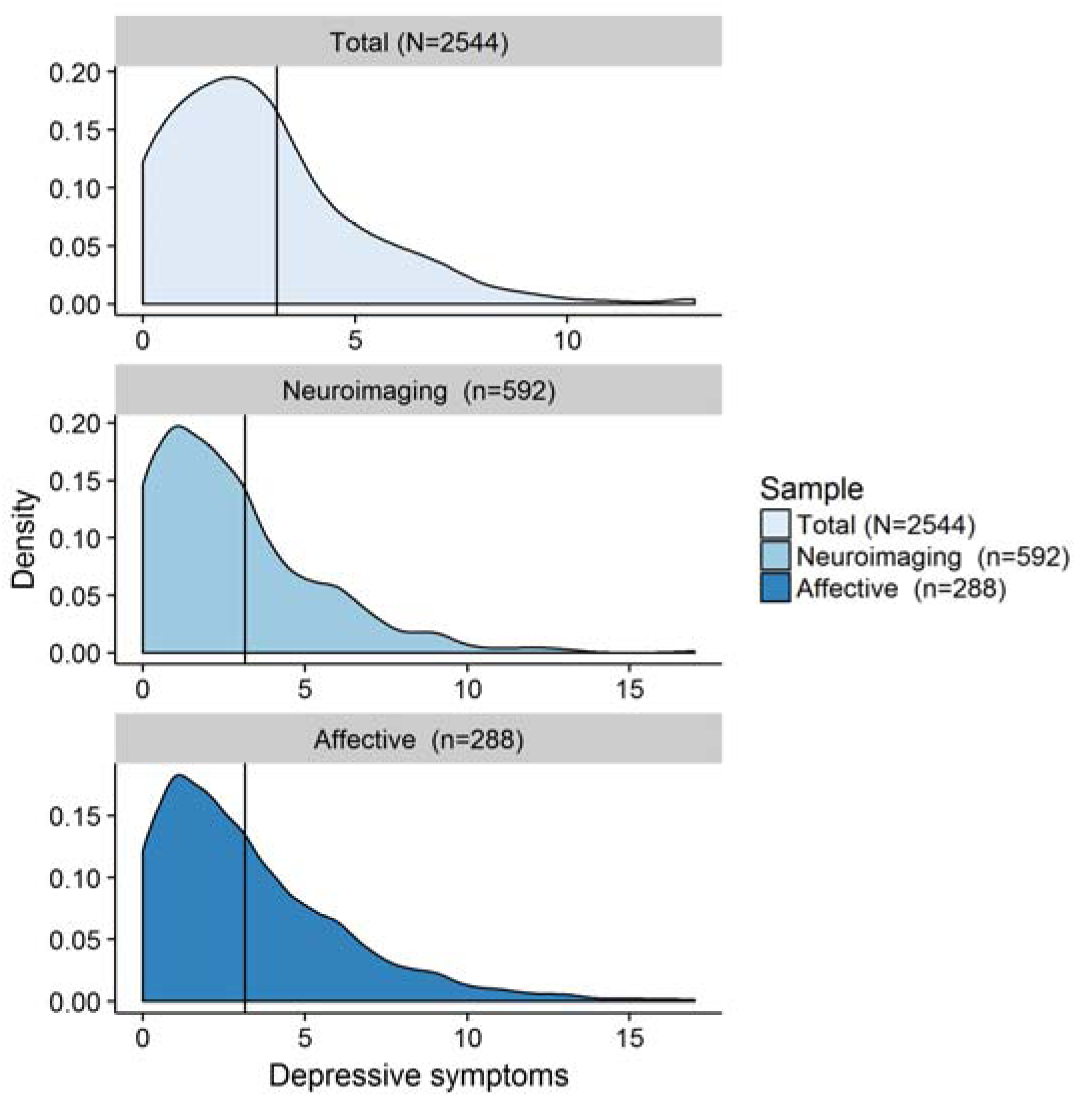
Neuroimaging cohort = individuals from the overall cohort for whom structural neuroimaging data is available; Affective cohort = individuals from the overall cohort who completed the valenced memory measure; *N/n* = number of participants; Depressive symptoms = number of self-reported symptoms of depression on the Hospital Anxiety and Depression Scale (HADS) depression sub-scale (range: 0-21 Zigmond and Snaith, 1983).

Importantly, the present study allowed us to investigate the relationship between these symptoms of depression and 1) subjective memory complaints, 2) objective memory on a standardized test, and 3) objective memory in a more specialized, affective context. Given the non-normal distribution of depressive symptoms in the cohort (Figure 1), we ran all analyses as non-parametric tests. More specifically, we entered the HADS-scores into a non-parametric logistic regression analyses based on ranks for the dichotomous outcome (i.e., subjective memory complaints) and non-parametric regression for the continuous outcomes (i.e., objective memory measures). Given the directionality of our hypotheses all significance testing of our *a priori* hypotheses was one-tailed, significance level for all exploratory tests were two-tailed.

In the full cohort (*N* = 2544), more symptoms of depression were related to more frequent self-report of memory complaints, *β* = 6.99^−4^, *SD* = 5.94^−5^, 95%CI [5.83^−4^; 8.16^−4^], *z* = 11.76, *p* ≤ .001, *R*^2^_*Nagelkerke*_ = .08 (H1a) and poorer performance on a standardized measure of memory *β* = −1.00^−3^, *SD* = 1.32^−4^, 95%CI [−1.00^−3^; −8.58^−4^], *t*(2542) = −8.44, *p* ≤ .001, *R*^2^ = .03 (H1b, Figure 2A). However, only the relationship between depressive symptoms and subjective memory survived adjustment for age, cognitive ability and gender^2^, *β* = 5.19^−4^, *SD* = 6.37^−5^, 95%CI [3.94^−4^; 6.44^−4^], *z* = 8.15, *p* ≤ .001, *R*^2^ _*Nagelkerke*_=.20; the same was not true for the relationship with standardized memory performance, *β*= −1.31^−4^, *SD* = 1.14^−4^, 95%CI [−1.00^−3^; −8.58^−4^], *t*(2539) = −1.15, *p* = .125, *R*^2^=.33, *R*^2^ _*adj*_=.33 (Figure 2B). The absence of a relationship was confirmed with Bayesian analysis (JASP Team, 2016), BF_01_ = 10.85. Moreover, exploratory analyses showed that the relationship between symptoms of depression and subjective memory complaints did not appear to be due to individuals who suffer from symptoms of depression simply reporting more neuropsychiatric health complaints. That is, while symptoms of anxiety were related to subjective memory complaints, *β* = 2.35^−4^, *SD* = 5.63^−5^, 95%CI [1.24^−4^; 3.45^−4^], *z* =4.16, *p* ≤ .001, *R*^2^ _*Nagelkerke*_ =.01, the relationship was no longer significant when accounting for depressive symptoms in the same analyses, *β* = −5.63^−5^, *SD* = 6.30^−5^,95%CI [−1.80^−4^; 6.71−4], *z* = −0.89, *p* =.372, *R*^2^ _*Nagelkerke*_ = .00, whereas the relationship between depression and subjective memory complaints remained significant after adjusting for symptoms of anxiety, *β*= 7.23^−4^, *SD* = 6.52^−5^, 95%CI [5.96^−4^; 8.51^−4^], *z* = 11.09, *p* ≤ .001, *R*^2^ _*Nagelkerke*_ =.08.

**Figure 2.**
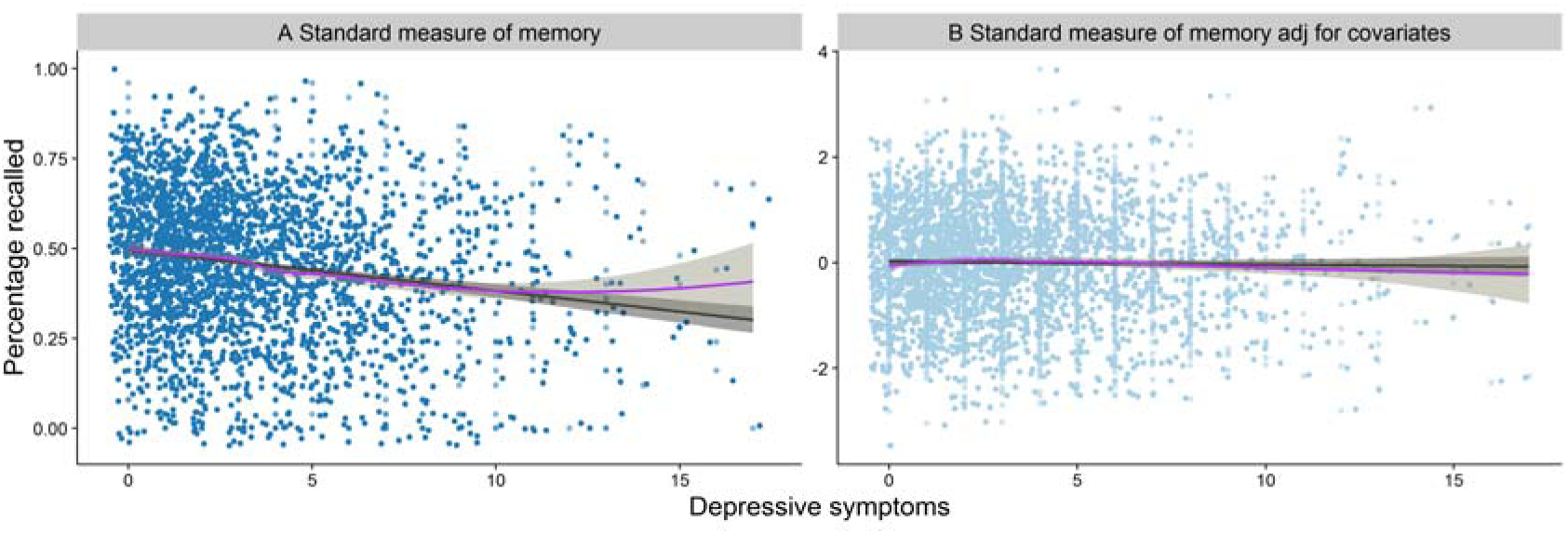
The figure represents the relationships between: A) depressive symptoms and performance on the standard measure of objective memory; B) 2A, after adjustment for age, cognitive ability and gender; jitter was added to the distribution for illustration purposes.

In the neuroimaging cohort (*n* = 592), multiple regression including total intracranial volume (TIV) as a covariate showed that hippocampal volume was related to both subjective memory, *β* = −4.64, *SD* = 1.71, 95%CI [−8.02; −1.23], *z* = −2.71, *p* =.004, *R*^2^ _*Nagelkerke*_ = .02, and the standardized objective measure, *β* = 17.94, *SD* = 3.29, 95%CI[11.47; 24.41], t(589) = 5.45, *p* ≤ .001, *R*^2^=.05, *R*^2^ _*adj*_ = .04 (Figure 3A). However, neither the relationship between hippocampal volume and subjective memory complaints, β = 0.43, *SD* = 2.00, 95%CI [−3.50; −4.36], *z* = 0.21, *p* = .416, *R*^2^ _*Nagelkerke*_ =.07, Nor relationship between hippocampal volume and objective memory *β* = 0.22, *SD* = 3.40,95%CI [−6.45; 6.90], *t*(586) = 0.07, *p* = .474, *R*^2^=.25, *R*^2^_*adj*_ = .24, survived adjustment for age, cognitive ability and gender. Contrary to our expectations, there was no significant association between depressive symptoms and hippocampal volume, *β* = −2.57, *SD* =2.01, 95%CI [−6.51; 1.38], *t*(589) = 1.28, *p* = .100, *R*^2^= .00(Figure 3B), which was again supported by a Bayes factor of 6.87 in favor of the null hypothesis. Unsurprisingly therefore, the relationships between depressive symptoms and both subjective *β* = 3.00^−1^, *SD* = 5.59^−4^, 95%CI [2.00^−1^; 4.00^−1^], *z* = 5.04, p ≤ .001, *R*^2^ _*Nagelkerke*_ = .08 and objective *β* =−2.00*−1*, *SD* = 1.00^*−1*^, 95%CI [−4.00^−1^; 1.03^−4^], *t*(588) = −1.86, *p* = .032, *R*^2^= .05, *R*^2^ _*adj*_ =.05 memory remained significant after adjusting for hippocampal volume (and TIV). That is, there was no support for the hypothesis that hippocampal volumes account in part for the relationship between depressive symptoms and memory performance in this non-clinical population (H2).

**Figure 3.**
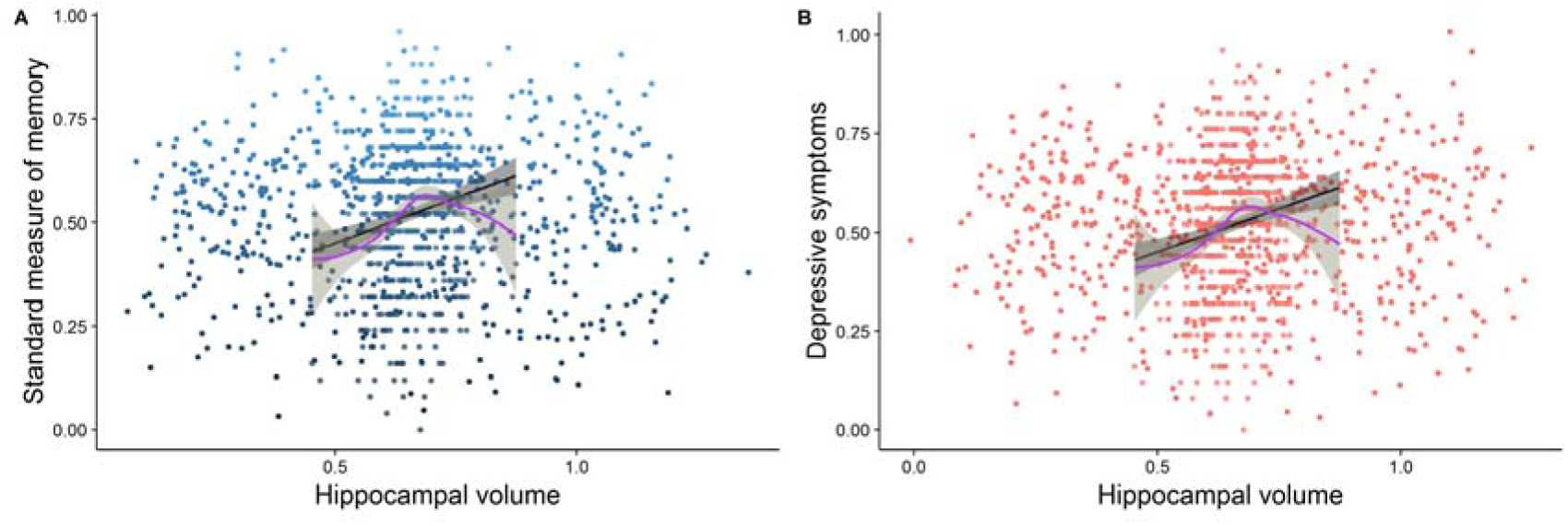
The figure represents the relationships between: A) hippocampal volume and performance on the standard measure of memory; B) hippocampal volume and symptoms of depression; jitter was added to the distribution for illustration purposes.

In the cohort that completed the measure of memory in affective contexts (*n* = 288), we investigated whether depressive symptoms showed a differential association with memory performance in affective contexts compared to a standardized measure of memory (H3). As in the overall sample, the relationship between depressive symptoms and performance on a standardized memory measure, *β* = −5.00^−1^, *SD* = 3.00^−1^, 95%CI [−1.10^−1^; 6.91^−4^], *t*(286) = −1.78, p = .043, *R*^2^ =.01, did not survive adjustment for age, cognitive ability, and gender, *β* = −2.00^−1^, *SD* = 3.00^−1^, 95%CI [−7.00^−1^; 3.00^−1^], *t*(283) =−0.66, *p* = .509, *R*^2^= .23, *R*^2^ _*adj*_ = .21, BF01 = 4.69. Symptoms of depression were, however, significantly related to poorer memory performance in negative, *β* = −4.00^−2^, *SD* = 1.00−2, 95%CI [−6.00^−2^; −1.00^−2^], *t*(286) = −3.48, *p* ≤ .001, *R*^2^ = .04 and positive *β* = −2.00^−2^, *SD* = 1.00^−2^, 95%CI [−5.00^−2^; −6.39^−5^], *t*(286) = −2.52, *p* = .006, R^2^ = .02contexts (Figure 4). However, when adjusting for performance in neutral contexts only the relationship between depressive symptoms and negative context remained significant, *β* = −1.45*−4*, *SD* = 5.93^−5^, 95%CI [−2.61^−4^; −2.79*−5*], *t*(285) = −2.44, *p* = .015, *R*^2^ = .76, R^2^ _*adj*_ = .76, even after adjusting for the same covariates, *β* = −1.38^−4^, *SD* = 5.78^−5^, 95%CI [−2.51^−4^; −2.40^−5^], *t* (285) = −2.38, *p* = .018, *R*^2^ = .78, *R*^2^ _*adj*_ = .78., but depressive symptoms were not related to memory in positive contexts after adjusting for performance in neutral contexts, *β* = −3.22^−5^, *SD* = 5.85^−5^, 95%CI [−1.47^−4^; 8.32^−5^], t(285) = −0.55, *p* = .584, *R*^2^ = .76, *R*^2^ _*adj*_ = .75 and covariates, *β* = −2.15^−5^, *SD* = 5.76^−5^, 95%CI [−1.00^−3^; 9.19^−5^], *t*(282) = −0.37, *p* = .709, *R*^2^ = .77, *R*^2^ _*adj*_ = .76.

**Figure 4.**
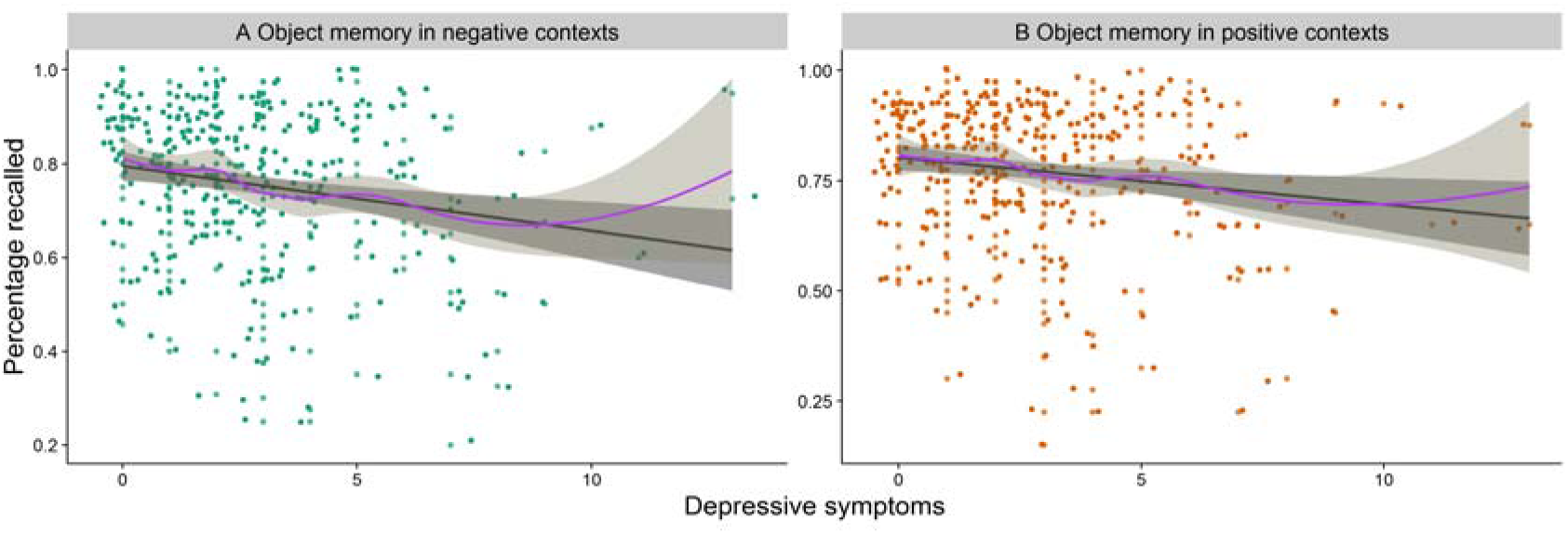
The figure represents the relationships between: A) depressive symptoms and memory for objects presented in negative contexts; B) depressive symptoms and memory for objects presented in positive contexts; jitter was added to the distribution for illustration purposes.

Furthermore, when directly comparing the size of the relationship of depressive symptoms with memory for objects in negative contexts with that for objects in positive contexts, the former was significantly higher, Williams’ *t*(285) = 3.07, *p* = .002. There was a marginal relationship between symptoms of depression and memory for objects presented in negative contexts to be stronger than their relationship with memory on a standardized measure of memory, Williams’ *t*(285) = −1.70, *p* = .045 (H3).

## Discussion

The present study examined the memory correlates of depressive symptoms in a large, population-derived cohort. First, we showed that depressive symptoms were related to self-reported memory problems, even after controlling for variations in age, cognitive ability, and gender. Moreover, the relationship was not simply due to individuals’ tendency to report more mental health problems, as the relationship between depressive symptoms and subjective memory remained after adjusting for symptoms of anxiety. One possibility is that the association between symptoms of depression and subjective memory reflects a negative interpretative bias. This notion, known as ‘depressive realism’, suggests that individuals who report symptoms of depression show less positivity bias (Mezulis et al., 2004; Watson et al., 2007). Future research should therefore investigate whether other types of self-reported cognitive functioning problems, for example attentional control problems (Derryberry and Reed, 2002), are also selectively associated with symptoms of depression but not other measures of mental health functioning.

A second set of findings showed that depressive symptoms were also related to performance on a standardized test of memory, but in this case, we could not rule out the possibility that this relationship was due to variations in memory as a function of age, cognitive ability, and/or gender (which were all significantly related to objective memory; see Table S2). In line with Fried and Kievit (Fried and Kievit, 2016), we also found no evidence for a significant relationship between depressive symptoms and hippocampal volume in this non-clinical cohort. Future research could investigate alternative sources of brain alterations associated with commonly experienced symptoms of depression (e.g., 52).

Finally, a third investigation showed that, while performance on a standard memory test may be unaffected in individuals experiencing subclinical symptoms of depression, objective memory impairments are found when the memoranda are encountered in negatively-valenced settings. More specifically, depressive symptoms were related to worse recognition memory for visual objects presented against negative backgrounds, even when adjusting for age, cognitive ability and gender. Importantly, this relationship remained even when further adjusting for recognition memory for the same types of objects presented against neutral backgrounds. This suggests that the relationship was specific to the valenced context, rather than differences between the visual object recognition memory test and the standardized verbal recall test, in terms of, for example, the nature of the memoranda or retrieval demands. Furthermore, the relationship between depressive symptoms and recognition memory for objects in positive contexts was no longer significant after the same adjustment for memory in neutral contexts, and the size of the relationship between depressive symptoms and recognition memory for objects in negative contexts was significantly greater than that between depressive symptoms and memory in positive contexts. In other words, the sensitivity of memory to depressive symptoms was selective to memory in negative contexts.

The implication of this finding is that measures of memory in negatively-valenced contexts (e.g., Henson et al., 2016) may be particularly sensitive to subtle differences in memory performance caused by current affective state. Again it remains an open question as to whether the impact of subclinical depressive symptoms in the general population islimited to memory performance in negative contexts, or whether this extends to other types of higher cognitive functions in negatively-laden environments. Future research should also investigate whether the strength of the association between individuals’ memory performance in negative contexts and their symptoms of depression has predictive utility for the development of more severe clinical forms of depressive disorders. That is, whether memory for neutral information in negative context fits within a larger pattern of cognitive vulnerabilities to depression (Gotlib and Joormann, 2010).

In conclusion, these findings show that the frequency of self-reported memory problems increases as a function of depressive symptoms. However, depressive symptoms are not associated with memory performance on a standard objective memory measure, when controlling for age, general cognitive ability and gender. Rather, depressive symptoms are associated with poorer memory for objects presented in negative contexts. These results suggest that memory for objects presented in negative contexts may be particularly sensitive to the memory problems reported by those experiencing symptoms of depression.

## Footnotes

1 We also ran all analyses excluding the 9% of participants who scored above the clinical cutoff (≥8), and found pattern of associations virtually unchanged

2 See Table S2 for the association between the memory measures and the covariates: age, cognitive ability and gender.

